# The p53 Protein is a Suppressor of Atox1 Copper Chaperon in Tumor Cells Under Genotoxic Effects

**DOI:** 10.1101/2023.07.25.550476

**Authors:** Sergey A. Tsymbal, Alexander G. Refeld, Viktor V. Zatsepin, Oleg A. Kuchur

## Abstract

The p53 protein is crucial for regulating cell survival and apoptosis in response to DNA damage. However, its influence on therapy effectiveness is controversial: when DNA damage is high p53 directs cells toward apoptosis, while under moderate genotoxic stress it saves the cells from death and promote DNA repair. Furthermore, these processes are influenced by the metabolism of transition metals, particularly copper since they serve as cofactors for critical enzymes. The metallochaperone Atox1 is under intensive study in this context because it serves as transcription factor allegedly mediating described effects of copper. Investigating the interaction between p53 and Atox1 could provide insights into tumor cell survival and potential therapeutic applications in oncology. This study explores the relationship between p53 and Atox1 in HCT116 and A549 cell lines with wild type and knockout TP53. The study found an inverse correlation between Atox1 and p53 at the transcriptional and translational levels in response to genotoxic stress. Atox1 expression decreased with increased p53 activity, while cells with inactive p53 had significantly higher levels of Atox1. Suppression of both genes increased apoptosis, while suppression of the ATOX1 gene prevented apoptosis even under the treatment with chemotherapeutic drugs. The findings suggest that Atox1 may act as one of key elements in promotion of cell cycle under DNA-damaging conditions, while p53 works as an antagonist by inhibiting Atox1. Understanding of this relationship could help identify potential targets in cell signaling pathways to enhance the effectiveness of antitumor therapy, especially in tumors with mutant or inactive p53.

## Introduction

With the accumulation of data on the antitumor effects of radio- and chemotherapy, numerous attempts have been made to identify the molecular mechanisms of survival and death of malignant cells. One of the most obvious markers, whose history began more than 40 years ago, is the oncosuppressor p53. This protein is a crucial regulator of tumor survival and death, an inducer of apoptosis and reparative processes, also plays an important role in cell response to ROS damage [1-4]. Furthermore, the balance of redox reactions in the cell is closely linked to the regulation of intracellular homeostasis of transition metals such as zinc (Zn), iron (Fe), and copper (Cu) [5-10]. However, there is limited information available regarding the correlation or codependence between the expression levels of p53 and proteins involved in metal metabolism in tumors [11-14]. Given the unique properties of copper and copper-binding proteins, investigating the metabolism of this metal becomes particularly attractive for developing approaches to combined tumor therapy [15, 16]. Copper plays a crucial role in redox reactions and the elimination of ROS as it is an integral part of the superoxide dismutase enzyme [17]. Additionally, copper can influence the level of intracellular glutathione, a major antioxidant molecule in cell [18]. Despite these important functions, the understanding of copper’s involvement in the occurrence and progression of tumor diseases is still in its early stages. Recent studies have focused on the dysregulation of copper-associated metallochaperones and enzymes during oncogenesis, as well as their potential therapeutic applications [19, 20]. Notably, enhancing the antitumor activity of disulfiram through the addition of copper ions has shown promising effects [21, 22]. Moreover, research conducted at the Laboratory of Diagnostics and Targeted Radiopharmaceutical Therapy of the University of Wisconsin has demonstrated a decrease in copper transport into the nucleus with inhibition or absence of p53 expression [23]. Further investigations have revealed a correlation between copper ion concentration and the activity of signaling cascades associated with malignancy, such as B-Raf, Akt, and HIF1 [24]. Inhibition of various copper carriers or chelation of copper ions also affects corresponding pro-oncogenic signaling pathways, including ERK, MAPK, NF-kB, and EGFR/Src/VEGF, which are involved in angiogenesis [25-28]. These findings suggest an association between p53 and copper-dependent proteins in tumor progression, highlighting the involvement of this tumor suppressor in the regulation of copper metabolism.

Considering our knowledge of the importance for oncotherapy of such copper-associated proteins as SOD1, CTR1, and angiogenin [5, 6, 29-31], an equally important player in copper metabolism, the Atox1 chaperone - which is an antioxidant and a transcription factor – remains aside. The role of this protein in tumor responses to genotoxic effects was unclear until recently. Only in 2015, a group of scientists from the Department of Hematology and Medical Oncology at Emory University showed that inhibition of Atox1 directly reduces the proliferation of tumor cells [32], and the binding of Atox1 to the cis-element of Cyclin D1 stimulates the growth and proliferation of mouse embryonic fibroblasts, as well as SW480 and SW620 colorectal cancer cells [33, 34]. Atox1 also shown to influence DNA repair by transcriptionally activating MDC1 protein [35]. Knockdown of Atox1 in non-small cell lung cancer cells reduces proliferative and growth processes [36]. Apparently, p53 activation, depending on the cell line and type of exposure, can differently affect the expression of Atox1, the induction of which protects the cell from death under ionizing radiation and cytotoxic drugs by eliminating ROS [37].

As a result, the data available in the literature on this topic are limited and rather contradictory. However, the general trend towards the study of copper metabolism and its relationship with typical cancer markers is very clear. We continue this trend, but our goal is to elucidate the role of the p53 tumor suppressor in the regulation of one particular participant in the copper metabolism pathways, Atox1, by paying attention to the responses of this protein to typical for tumor therapy stimuli, such as cytotoxic drugs and ionizing radiation. The data will lay the foundation for further research on this topic with the possible implementation of the development of new anticancer drugs.

## Results

### Atox1 activity is increased in cells with TP53^-/-^ gene

At the first stage, we assessed the basic level of gene expression and induction of the Atox1 protein in HCT116 colorectal cancer and A549 lung carcinoma cell lines with the wild type (WT) or inactivated by the CRISPR-Cas9 tumor suppressor gene TP53 (TP53^-/-^). Immunoblotting analysis revealed that cells with functional p53 exhibited reduced Atox1 activity, whereas p53 knockout cells showed a significant increase in Atox1 protein content by approximately 2.2-2.8 times under normal conditions. This trend was observed in both cell lines, with no statistically significant difference in the baseline levels of Atox1 protein between the two lines (Fig. 1A). To further validate our findings, we used p21, a p53-dependent inhibitor of cyclin-dependent kinase 1, as an additional control. Accumulated p53 activates the CDKN1A gene, leading to cell cycle arrest at the G_2_/M phase and inhibition of Cdc25, thereby facilitating DNA repair processes [38]. Consequently, the level of p21 decreases when p53 is suppressed. In p53 knockout cells, the amount of p21 protein was found to be reduced by approximately 2-fold compared to the control.

**Figure 1.**
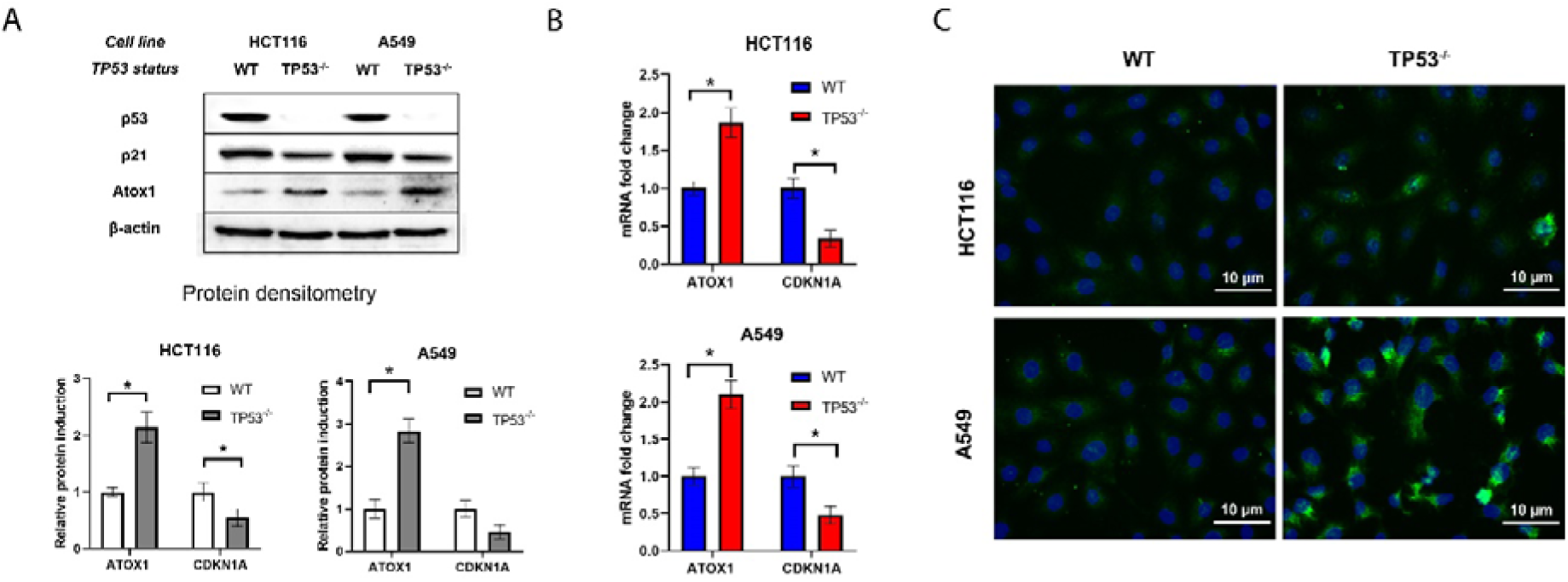
Dependence of Atox1 and p53 level in HCT116 and A549 cell lines with different TP53 status: **A** - immunoblotting with antibodies to p53, p21, and Atox1; β-actin was used as a normalization. A densitometric analysis of the obtained data is shown below. **B** – RT-qPCR analysis with primers for TP53, CDKN1A, and ATOX1 genes; GAPDH gene was used as a reference. **C** - immunofluorescence staining with primary antibodies to Atox1, secondary antibodies with AlexaFluor488. DAPI was used for nuclei staining. WT – wild type cells, TP53^-/-^ – cells without TP53. For all experiments: n = 3, mean +/− SEM, paired Student t-test, p < 0,05.

We also examined the transcriptional regulation of Atox1 gene in relation to TP53 status. Real-time PCR analysis revealed that the relative expression of Atox1 mRNA was approximately 2-2.5 times higher in p53 knockout cells compared to wild-type cells (normalized to 1.0), while the expression of CDKN1A, which encodes p21, decreased by approximately 2-fold (Fig. 1B).

Furthermore, immunofluorescence microscopy allowed us to visualize the observed pattern of increased Atox1 activity in cells with TP53 inactivation. The A549 cell line exhibited approximately 2.5 times higher levels of the metallochaperone compared to the HCT116 cell line (Fig. 1C). However, this method did not enable the detection of Atox1 translocation into the nucleus as a transcription factor, as reported in previous studies [37].

These findings raise important questions regarding the role of Atox1 as a p53-dependent factor, which is known to be a crucial sensor for responses to DNA damage, cell cycle regulation, and repair processes. Specifically, it prompts us to investigate whether Atox1 activity is altered in response to cytostatic and cytotoxic effects and whether it contributes to the regulation of the survival-death balance in tumor cells, particularly those harboring p53-null mutations.

### Atox1 is induced in a p53-dependent manner during genotoxic stress

According to Beaino W. et al., Atox1 protein is induced in a p53-dependent manner in response to the cytotoxic drug cisplatin [37]. According to the authors, this can be explained by the ability of cisplatin to a certain extent to replace copper ions and bind to Atox1, acting as a cofactor, which induces this protein. However, given the ability of Atox1 to act as a transcription factor and play a role in the processes of response to external stimuli [33], we formulated two hypotheses: 1) expression of Atox1 increases in response to various genotoxic signals (cytotoxic drugs, ROS inducers, ionizing radiation) at the level of transcription and translation; 2) this induction is p53-dependent.

Let’s start with the second: Indeed, as shown above and will be discussed below, Atox1 expression is increased in TP53 knockout sublines (HCT116TP53^-/-^, A549TP53^-/-^), which contradicts previous work by Beaino et al., who observed Atox1 induction only in HCT116 WT, while in cells with TP53^-/-^ the amount of Atox1 decreased by almost 2 times [37]. In this study, the effect was not replicated in the MEF mouse fibroblast line. On the contrary, we show a sequential pattern of Atox1 activation in HCT116 and A549 cell lines with inactivated TP53 both transcriptionally and at the translational level.

Returning to the first hypothesis, ATOX1 expression is clearly upregulated in response to multiple genotoxic stimuli. Thus, p53 reacts similarly to all genotoxic agents, except for hydrogen peroxide, both at the mRNA and protein levels, which is quite expected given its role in the response to DNA damage [39], which correlated with the results of our experiments (Fig. 2). At the same time, in the HCT116TP53^-/-^ and A549TP53^-/-^ cell lines, Atox1 is activated upon the addition of 0.1μM doxorubicin, 80nM PMA, and 10μM bleomycin (and, to a certain extent, 35μM cisplatin in wild-type cells) and is weakly activated when p53 is normally functioning. Thus, in HCT116 cells with suppressed TP53, Atox1 induction in response to cisplatin, doxorubicin, PMA, and bleomycin was 2.1, 2.5, 3.0, and 2.8 times higher compared to the control, respectively. For the A549TP53^-/-^ line, these values were 2.0, 2.8, 2.4, and 2.2 times, respectively (Fig. 2A). The weak response to H_2_O_2_ is apparently associated with its short half-life/rapid decay and rapid cellular responses that regulate changes in the cell in response to oxidative stress, which does not lead to large-scale translational responses. It is worth noting that Atox1 inducibility is on average similar for both cell lines to the respective drugs, which was not observed in previous studies. Doxorubicin, PMA, and bleomycin caused the strongest differences in Atox1 expression in both lines; they were selected by us for PCR analysis.

**Figure 2.**
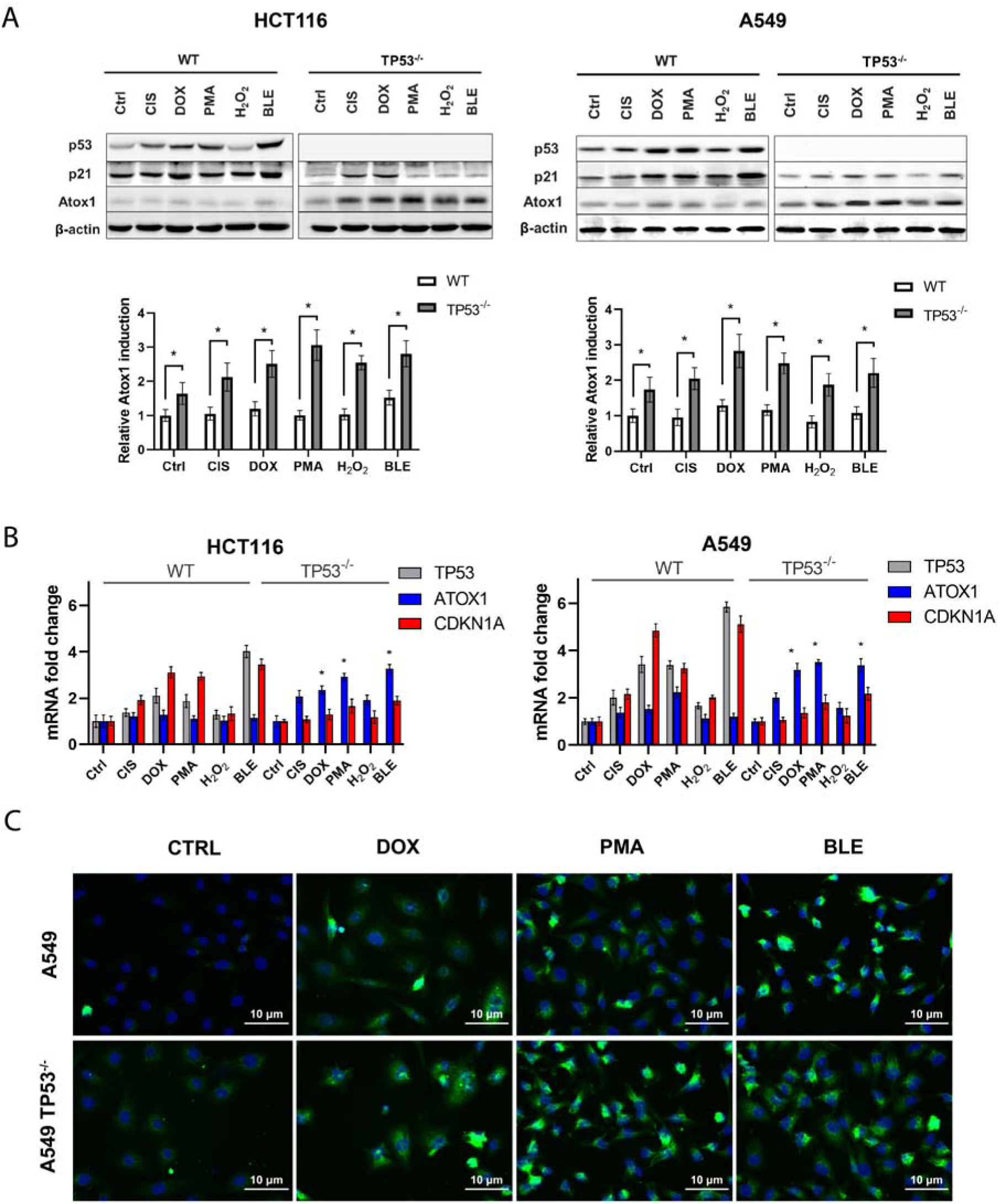
Influence of cytotoxic agents on the activity of Atox1 at different status (WT and KO) of the TP53 gene in A549 and HCT116 cell lines, 24h after drugs exposure. **A** - immunoblotting with antibodies to p53, p21, and Atox1; β-actin was used as a normalization. A densitometric analysis of the obtained data is shown below. **B** – RT-qPCR analysis with primers for TP53, CDKN1A and ATOX1 genes; GAPDH gene was used as a reference, **C** - immunofluorescence staining with primary antibodies to Atox1, secondary antibodies with AlexaFluor488. DAPI was used for nuclei staining. DOX – doxorubicin (0,1μM), CIS – cisplatin (35μM), PMA – phorbol-12-myristate- 13-acetate (80nM), H_2_O_2_ – hydrogen peroxide (450μM), BLE – bleomycin (10μM). WT – wild type cells, TP53^-/-^ – cells without TP53. For all experiments: n = 3, mean +/− SEM, two-way ANOVA, p < 0,05.

Expression analysis of the TP53, CDKN1A, and ATOX1 genes upon exposure to previously selected drugs confirmed the data obtained by immunoblotting (Fig. 2B). The control (intact cells, no effects) was taken as 1.0, the addition of doxorubicin to HCT116 and A549 cells led to an increase in the expression of TP53 and CDKN1A by 3-4 and 4-5 times, respectively, relative to the control, for PMA these values were equal to 3-4 for both genes relative to the control, respectively, for bleomycin - 5-6 in both cases. At the same time, Atox1 expression for all compounds did not exceed 2 change fold in the studied cell lines with wild type p53. In the case of TP53 knockouts, the CDKN1A gene was practically not expressed, and Atox1 activity increased to ∼3.5 change fold when exposed to doxorubicin and PMA (both cell lines), and up to 4.5 when bleomycin was added (Fig. 2B). In general, transcriptional and translational response data for chemotherapeutic agents were comparable.

It is generally believed that the main role of Atox1 lies in its functions as a transcription factor under stressful conditions, it also has a nuclear localization sequence [40]. Atox1 has been shown to migrate into the nucleus under the action of cisplatin in a p53-dependent manner [37]. To assess the nuclear translocation of Atox1, immunocytochemical staining of the Atox1 protein was performed under the influence of doxorubicin (0.1μM), PMA (80nM), and bleomycin (10μM), which showed good results in transcriptional and translational activation. Fluorescence microscopy showed no discernible nuclear translocation of Atox1 upon genotoxic exposure, with the protein increasing markedly, especially with doxorubicin and bleomycin (Fig. 2C). It is likely that the drugs used in this experiment, unlike cisplatin, do not have the ability to bind Atox1 and induce its migration into the nucleus, since they do not interact with copper metabolism proteins [41]. In addition, it has not previously been shown to increase Atox1 with PMA supplementation, but Atox1 is known to induce NFkB in inflammation [42-44].

In addition to cytotoxic drugs exposure, we investigated the impact of ionizing radiation on the transcriptional and translational responses of Atox1. Ionizing radiation is known to cause single- and double-strand DNA breaks and generate reactive oxygen species through water radiolysis [45, 46]. To generate gamma radiation, we utilized the RUM-17 radiotherapy unit, with an effective therapeutic dose of 10 Gray (Gy).

Our findings corroborate previous observations on the response of p53 to radiation. Specifically, in HCT116 cells, p53 activity increased by three-fold compared to the non-irradiated control, and in A549 cells, it increased by 3.6-fold. Similarly, the induction of the p21 protein followed a similar pattern, albeit with lower activity levels. The expression of p21 was 1.5-2 times higher than the control values, and its induction was reduced in cell lines with inactive TP53 but increased upon irradiation.

In contrast, the metal chaperone Atox1 exhibited minimal response to gamma radiation, irrespective of the p53 status. However, in irradiated A549TP53^-/-^ cells, there was a slight suppression of Atox1 induction compared to the same subline without irradiation (Fig. 3A).

**Figure 3.**
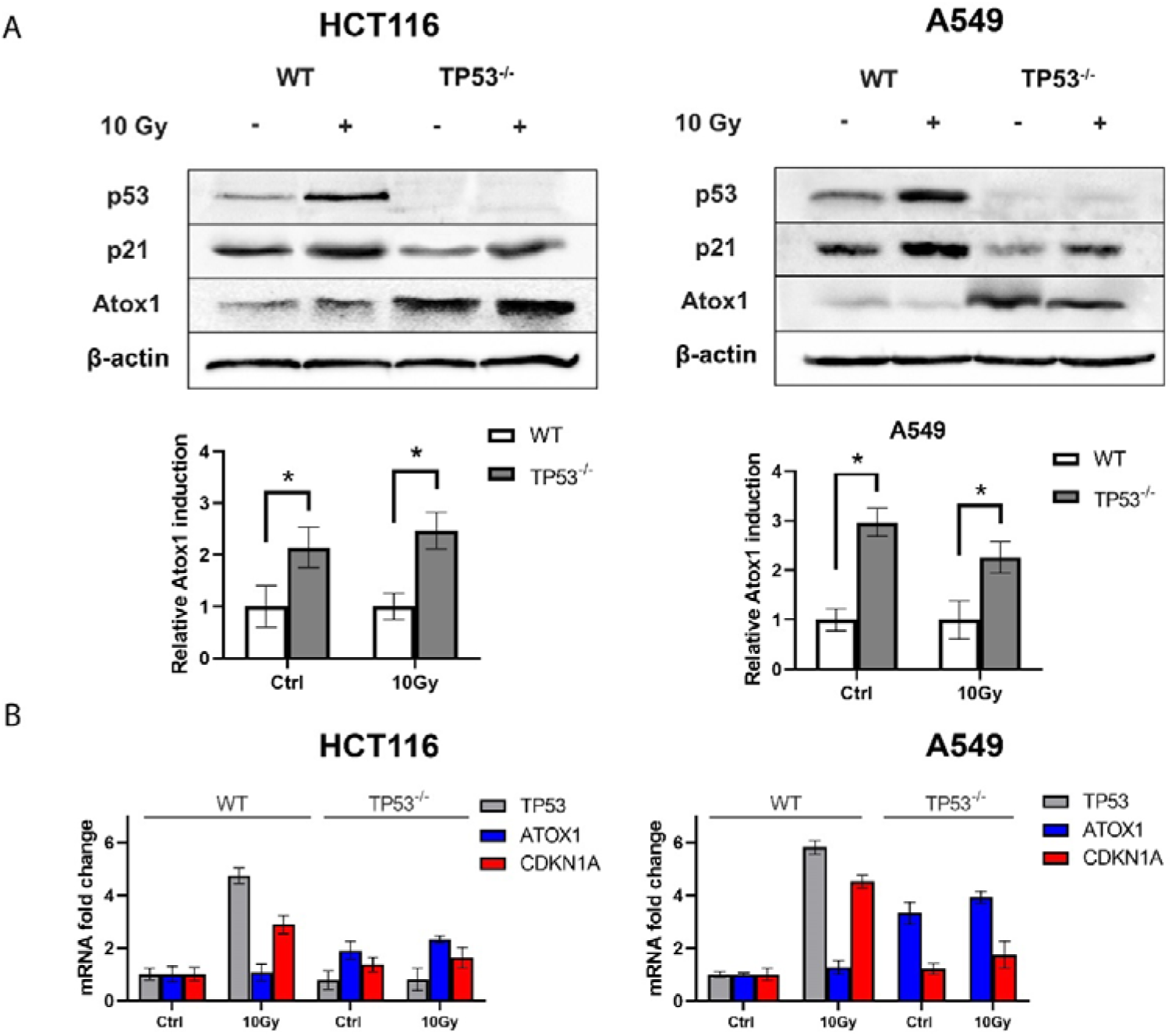
Influence of ionizing radiation on the activity of Atox1 at different status (WT and KO) of the TP53 gene in A549 and HCT116 cell lines, 24h after ionizing irradiation (10Gy) exposure. **A** - immunoblotting with antibodies to p53, p21, and Atox1; β-actin was used as a normalization. A densitometric analysis of the obtained data is shown below. **B** – RT-qPCR analysis with primers for TP53, CDKN1A and ATOX1 genes; GAPDH gene was used as a reference. The value of WT 0Gy (control) was taken as a 1.0 for all genes and is not shown in the graphs. For all experiments: n = 3, mean +/− SEM, two-way ANOVA, p < 0,05.

To validate these findings at the transcriptional level, we performed real-time PCR. Irradiation with a dose of 10 Gy resulted in a four to 6-fold increase in TP53 gene expression relative to the control. Conversely, Atox1 exhibited weak expression levels. Interestingly, the absence of TP53 led to the activation of Atox1, and radiation further enhanced this effect, particularly in HCT116 cells with TP53^-/-^, where Atox1 expression was approximately 3-4 times higher than in the intact control (Fig. 3B). Fluorescent microscopy with the distribution of Atox1 protein after exposure to ionizing radiation at a dose of 10 Gy are shown in the Supplementary Material (Figure S1)

The next experiments showed that in the absence of p53 the Atox1 protein can be induced by DNA-damaging agents (doxorubicin and bleomycin) but respond poorly to ROS exposure (H_2_O_2_, ionizing radiation). This effect is observed both at the transcriptional and translational levels. In addition, the Atox1 induction caused by the activation of NFkB by the addition of PMA is important observation, since only reverse regulation has been described so far. The specific role of Atox1 in response to these stimuli, as well as participation in the regulation of survival and adaptation processes, remains to be established. In our next experiments, we used the siRNA transient gene knockdown to identify the effect of inactivation of genes of interest on cell survival and response to genotoxic stress.

### The Influence of p53 on Atox1 Activity is Unidirectional

Cell culture conditions can induce significant changes in cell metabolism and gene expression upon permanent gene inactivation, impacting cell cycle regulation and viability [47]. To avoid these specific changes, we utilized siRNA-mediated knockdown or small molecule inhibitors for transient gene inactivation, allowing us to study the immediate effects of ATOX1, TP53, and their co-inactivation on their reciprocal regulation, as well as changes in cell viability and cell cycle.

To assess knockdown efficiency, we measured TP53 and ATOX1 expression levels using RT-qPCR. Our results demonstrated a 10-fold decrease in TP53 expression and a 100-fold decrease in ATOX1 expression (Fig. 4A). While we previously discussed p53-dependent changes in Atox1 levels, it remained unclear whether Atox1 directly influences p53 activity. To address this, we evaluated the reciprocal regulation of p53 and Atox1 proteins in cells with transient suppression of these genes. Western blot analysis revealed that, similar to HCT116TP53^-/-^ and A549TP53^-/-^ lines, the absence of functional p53 led to increased Atox1 activation (Fig. 2). However, Atox1 inactivation did not affect the levels of p53 or p21 (Fig. 4B). Simultaneous suppression of both genes (doubleKD) significantly reduced their expression, while some residual Atox1 was observed (∼10-20% of control). These dependencies were further enhanced with the addition of bleomycin (Fig. 4B). Similar results with the suppression of the TP53 and ATOX1 genes were shown by immunoblotting on the HTC116 cell line. Data is shown in the Supplementary Material (Figure S2)

**Figure 4.**
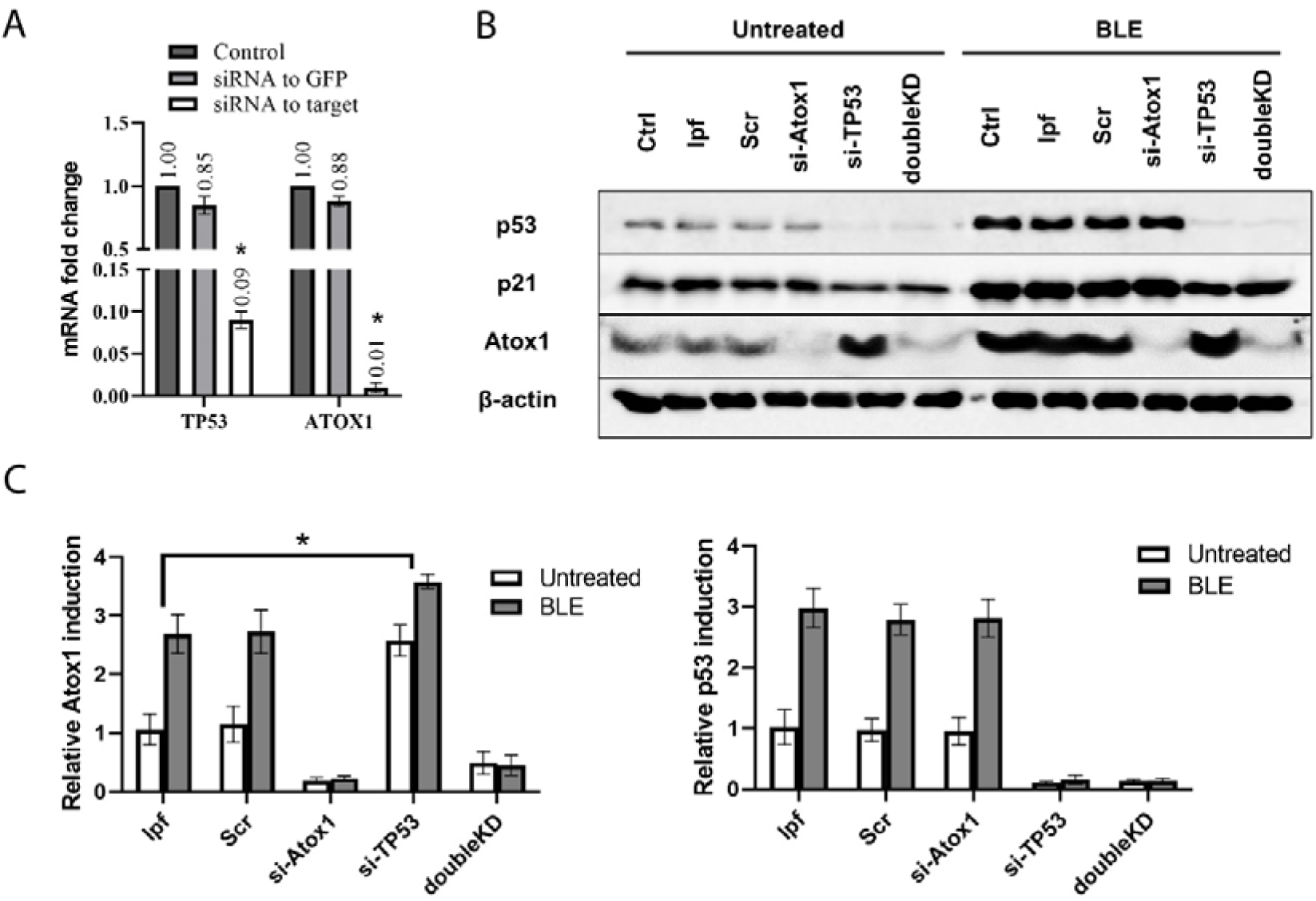
Analyzing the effect of ATOX1 and TP53 gene siRNA-mediated knockdown on mutual expression, A549 cell line. **A** - RT-qPCR analysis with primers for TP53 and ATOX1 genes after their siRNA knockdown, 24h after inactivation; GAPDH gene was used as a reference. **B** – immunoblotting with antibodies to p53, p21, and Atox1; β-actin was used as a normalization. ATOX1 (si-ATOX1), TP53 (si-TP53) or double ATOX1/ TP53 (doubleKD) knockdowns were used in the absence (Untreated) and presence (10μM BLE) of bleomycin, 24h after exposure. **C -** A densitometric analysis of the obtained data is shown below. Controls are taken as 1.0 and not shown on densitometry. For all experiments: n = 3, mean +/− SEM, two-way ANOVA, p < 0,05.

### Suppression of ATOX1 under genotoxic stress increases tumor viability, but simultaneously suppression of TP53 decreases it

The MTT assay on A549 cells made it possible to assess the viability of cells with active and inactive ATOX1 or TP53, when exposed to bleomycin or gamma radiation. The test showed that knockdowns by themselves did not affect cell viability under intact conditions, only double inactivation of the ATOX1 and TP53 genes (doubleKD) led to a ∼10–12% decrease in survival. On the first day (24 hours) of genotoxic effects, there are also no pronounced changes in cell survival. The addition of the genotoxic drug 10μM BLE or exposure to 10Gy of gamma radiation increased cell death after 72 hours: in the case of ionizing radiation, the survival rate decreased by 35% compare to the control, in the case of exposure to bleomycin, by 40%. The same is true for samples with lipofectamine (lpf) and scrambled siRNA to GFP (Scr). Further, it was found that the frequency of cell death with knockdown of the ATOX1 gene was reduced on the 3rd day after the respective treatments. Thus, the percentage of surviving ATOX1-negative cells after 72 hours after they were exposed to gamma radiation and treatment with bleomycin was 86% and 84.5%, respectively. The control values for wild-type cells after the respective treatments were 72.4% and 67.2%, respectively. At the same time, knockdown of TP53 reduced cell viability compared to the control: when exposed to radiation and bleomycin, the survival of cells with inactivated TP53 after 72 hours was 60.9% and 49.2%, respectively. Finally, double gene knockdown resulted in marked cell death: 37.6% and 31.9% on radiation and bleomycin exposure, respectively (Fig. 5A). Thus, ATOX1 inactivation serves as a kind of “protector” of cells from death, but with simultaneous inactivation of TP53, this property is also removed, and the opposite effect is observed - inhibition of cell survival.

**Figure 5.**
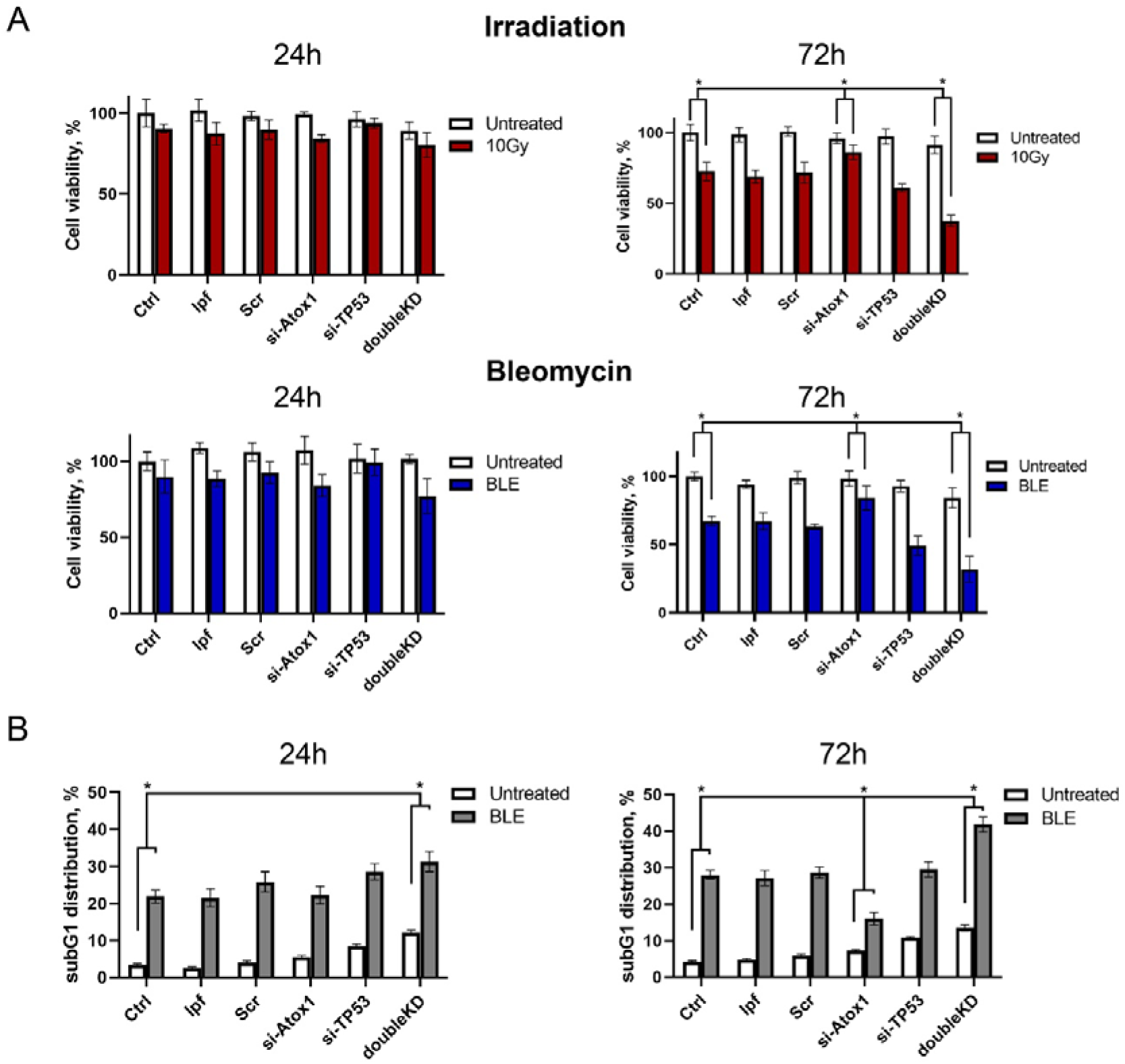
Influence of ATOX1, TP53, and double knockdown on cell viability in the absence and presence of genotoxic stress in A549 cell line. **A** – MTT-assay after ATOX1 (siATOX1), TP53 (siTP53) or double ATOX1/ TP53 (doubleKD) knockdowns in the absence (Untreated) and presence of ionizing irradiation (10Gy) and 10μM bleomycin (BLE) at the different time points: 24 or 72 hours after exposure. **B** – subG_1_ phase accumulation caused by TP53 (siTP53), ATOX1 (siATOX1) or double genes (doubleKD) knockdowns in the absence (Untreated) and presence of 10μM bleomycin (BLE). For all experiments: n = 3, mean +/− SEM, two-way ANOVA, p < 0,05.

To elucidate the reasons for the observed effects of death avoidance upon inactivation of ATOX1, we examined the distribution of cell cycle phases using A549 cell line (as the line that most effectively responds to stressful conditions at the transcriptional level) under the same conditions (TP53, ATOX1 or knockdown of both genes, with or without bleomycin). The addition of siRNA to TP53 and ATOX1 in the case of untreated cells (addition of 250 nM lepofectamine 2000) practically did not change the distribution of G_1_, S and G_2_/M phases after 24-72 hours (subG_1_ < 10%), a different situation was observed with simultaneous knockdown of TP53 and ATOX1 (doubleKD): while in the control group the subG_1_ phase was 2-5%, the absence of both genes led to an increase in this phase to 10-13%, while there were no noticeable changes in other phases, G1 and G_2_/M.

In the bleomycin-supplemented group (250 nM lpf 2000, 10 µM BLE), the differences were more pronounced (Fig. 5B). For example, 24 hours after the addition of bleomycin, about 20% of the cells are in the SubG_1_ phase, while after 72 hours the relative number of events in this phase rises to 29%. The addition of GFP siRNA (Scr) did not change the ratio between cell cycle phases relative to the controls described above. Suppression of TP53 and, accordingly, its reparative functions and control of cell cycle arrest, did not lead to an obvious increase in subG_1_: 18.2% at 24 hours after the addition of bleomycin and 25.5% at 72 hours, respectively. However, at this point, a time-dependent increase in the G_2_/M phase was observed (23.5% and 40.5%, respectively). Unexpected, but consistent with the logic of the MTT test, changes in the cell cycle were observed when ATOX1 was suppressed: the transition of cells to the subG_1_ fraction slowed down. So, 24 hours after the addition of bleomycin, subG_1_ (in the group with si-Atox1) was equal to 22% with values in the control group (lpf) 17%. However, after 72 hours, subG_1_ (si-Atox1) was 16% with values in the control group (lpf) of 28%. Finally, in the BLE group with inactivation of both genes (doubleKD), increased tumor cell death is observed with almost complete escape of cells from the G_2_/M phase. For example, 24 hours after the addition of bleomycin, the subG_1_ and G_2_/M phases were 31.3% and 18.8%, respectively. After 72 hours, these figures were 42% and 12.5%, respectively. According to our results, active Atox1 in cells with DNA damage can induce apoptosis, but the absence of its functioning form creates a block at the G_1_/S checkpoint and limits the ability of cells to go into apoptosis (subG_1_). Disabling the second gene, TP53, allows cells to bypass this effect and successfully redistribute into the subG_1_ phase.

## Discussion

In this study, we established that the expression of the transcription factor and antioxidant protein Atox1 is more pronounced in cell lines with inactivated TP53. Specifically, we observed that cell lines with wild type p53, such as HCT116 and A549, exhibited reduced gene expression and induction of Atox1 protein compare to cells with inactive p53, either through knockout or knockdown techniques. Furthermore, common antitumor drugs such as doxorubicin and bleomycin, as well as exposure to therapeutic doses of ionizing radiation, activated p53 in normal cells, while the concentration of Atox1 protein is significantly increased only in cells with inactive TP53. Conversely, the suppression of the ATOX1 gene using small interfering RNAs (siRNAs) did not result in any transcriptional or translational changes in the level of TP53 in these cell lines. These results suggest that p53 tumor suppressor acts as a negative regulator of Atox1, while no reciprocal feedback mechanism was identified. Fluorescent microscopy with antibodies to Atox1 revealed that presence of genotoxic stimulus caused only minor translocation and accumulation of Atox1 in the nucleus, suggesting permanent intranuclear localization of this protein, where it can possibly function as a transcription factor.

Moreover, our study examined the impact of siRNA-mediated knockdowns of the TP53 and ATOX1 genes on cell survival and cell cycle distribution in the absence of cytotoxic drugs. It was found that knockdown of these genes did not alter these parameters. When treated with the bleomycin, both cell lines exhibited an increased accumulation of cells in the subG_1_ phase (indicative of cell death), which was further enhanced with TP53 inactivation through siRNA knockdown. Surprisingly, we observed a decrease in the subG_1_ phase accumulation when bleomycin was added to cells with inactivated ATOX1. These findings were corroborated by the MTT assay. Lastly, we investigated the simultaneous knockdown of both ATOX1 and TP53 genes, which resulted in an increased apoptosis rate compared to cells with inactive TP53 alone. This effect was approximately two-fold higher 72 hours after drug exposure, while the number of G_2_/M-arrested cells decreased. This intriguing observation presents a paradoxical scenario, whereby inactivation of ATOX1 protects cells from death, but additional suppression of TP53 enhances the apoptotic effect by abolishing the G_2_/M cell cycle block and promoting cell death in the subG_1_ phase.

Collectively, these findings suggest the existence of a potential mechanism by which ATOX1 is inversely associated with p53 levels and facilitates cell death rather than, as previously proposed, cell survival by eliminating ROS [32, 33, 36, 37]. Moreover, in our experiments we did not observe significant increase of Atox1 level in response to H_2_O_2_ treatment, suggesting the primal role of Atox1 as a transcription factor, however, underlying cause for this phenomenon warrants investigation. While the role of p53 in governing the balance between repair and apoptosis in tumor cells has been extensively studied over the course of more than four decades [48-50], the precise involvement of Atox1 in this context remains enigmatic. Several evidences indicate that Atox1 can positively regulate the expression of cyclin D1, a key factor in cell cycle progression and the transition from the G_1_ to S phase [33, 37, 51]. It is plausible to speculate that inhibiting Atox1 upon exposure to DNA-damaging agents, such as chemotherapy or radiotherapy, prevents cells from bypassing the G_1_/S checkpoint with genomic damage (Fig. 6A). In the absence of Atox1-mediated CCND1 expression, cells fail to accumulate critical damage during DNA replication and do not undergo mitotic catastrophe [52]. This leads to a reduction in the subG_1_ and G_2_/M phases in ATOX1 knockdown experiments (Fig. 6B). Moreover, experimental data also elucidate the role of p53 as a negative regulator of Atox1: in TP53-inactivated cells, Atox1 expression is elevated, resulting in tumor cell death due to bypassing the G_1_/S and G_2_/M checkpoints without proper DNA repair (Fig. 6C).

**Figure 6.**
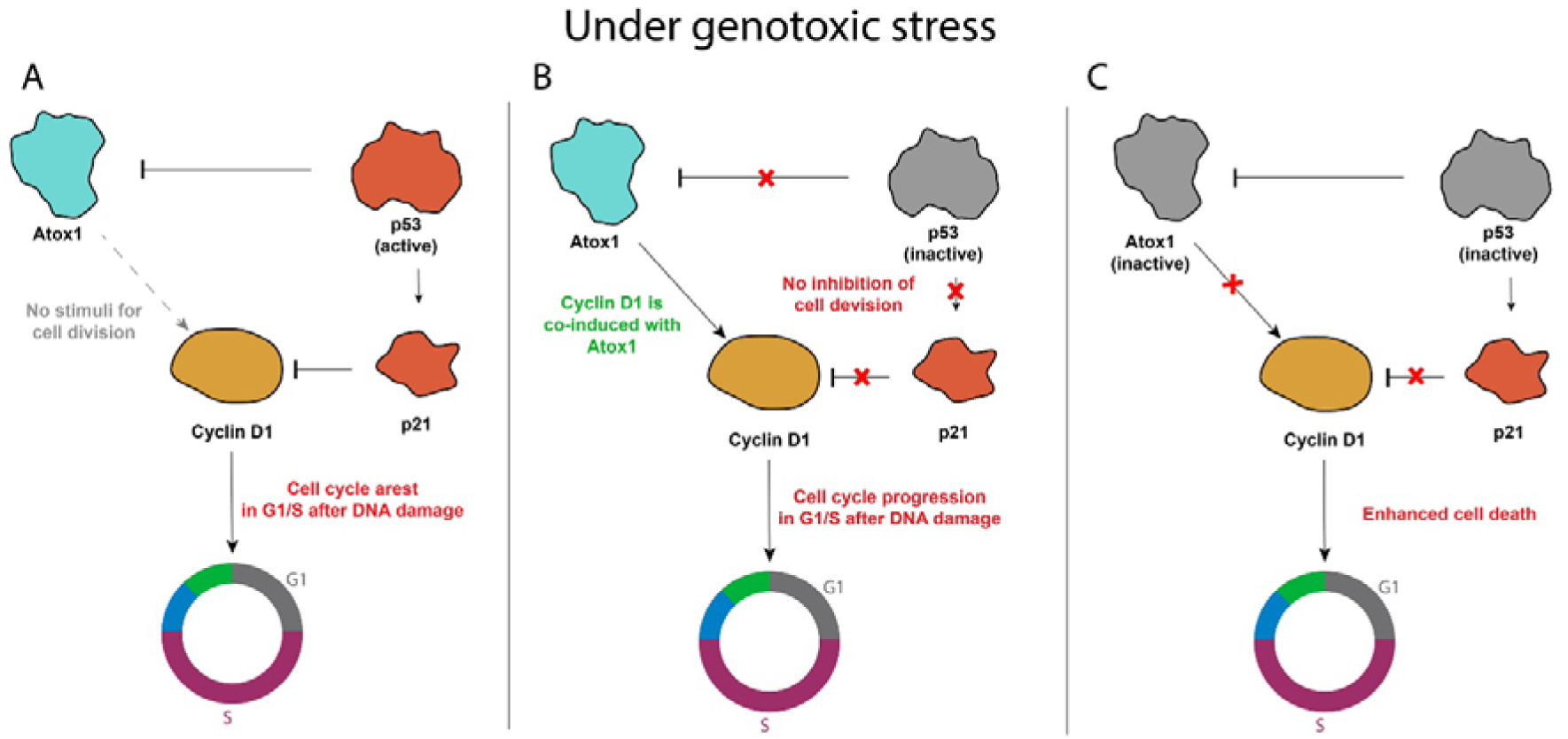
Hypothetical concept of Atox1-mediated cell cycle regulation via cyclin D1 and p53 influence on the process under genotoxic stress: **A** – when p53 fully functional Atox1 is suppressed as well as cyclin D1 (via p21 activation), **B** – when p53 is inactivated, Atox1 is released from suppression and promotes cell cycle progression due to the induction of cyclin D1, **C** – In case of double knockdown, when both Atox1 and p53 are suppressed, cells cannot properly regulate the transition from one phase of the cell cycle to another, resulting in increased sensitivity to DNA damaging agents

Nevertheless, this theory does not fully explain the synergistic impact of ATOX1/TP53 double knockdown. For instance, if the translocation of Atox1 into the nucleus was not conclusively observed during the experiment, how does it regulate cyclin D1? Is Atox1 involved in proapoptotic signaling, the inhibition of which protects cells from death? Could this be linked to the purported ability of Atox1 to function as a non-canonical modulator for the MAPK cascade, specifically mediating the phosphorylation of the transcription factor Erk (Ras-ERK signaling pathway, MAPK/ERK) [27, 53-55]? In this scenario, the p53-controlled Atox1-mediated regulatory network involving CCND1 may be even more intricate and context-dependent. Recent work about cuproptosis describes p53 participation in the regulation of this process [56] which may indicate that copper-mediated cell death possibly realized via Atox1. This could point broader Atox1 functions in cellular signaling related to cell survival. In any case, further investigation is warranted to unravel the role of Atox1-p53 in the regulation of Cyclin D1.

## Conclusion

In conclusion, present research lays the foundation for establishing a comprehensive and coherent framework for understanding the relationship between copper metabolism proteins and p53 activity in cell malignancy. This research opens the door for future studies to explore the role of Atox1 and p53 interaction in tumor progression and potential approaches to cancer therapy by targeting these proteins. Future investigations should focus on elucidating the underlying mechanisms of this interaction, including the involvement of Cyclin D1, p63 and p73, Ras-ERK and other proteins that regulate the G_1_/S and G_2_/M transitions. The identification of these factors and their association with p53 holds great promise for both diagnostic and therapeutic purposes. A deep and detailed study of these interactions in tumors of different localization under the influence of antitumor drugs and ionizing radiation agents will allow the development of optimal complex schemes for the treatment of tumors.

## Materials and Methods

### Cell lines and reagents

Transformed human cell lines used: HCT116 (colon adenocarcinoma) with intact p53; HCT116p53^-/-^ with a deletion of both alleles of the TP53 genes, as well as the A549 line with wild (A549) and knockout p53 (A549p53^-/-^) by the CRISPR-Cas9 method, acquired at ATCC. The cells were cultured in Dulbecco’s modified Eagle’s medium (DMEM, Biolot, Russia) supplemented with 2 mM L-glutamine, 10% fetal bovine serum (PAA, USA), and 100 U/mL gentamicin (Biolot). Only cells in the logarithmic growth phase, with no more than 15 passages, were used in the experiments. All other reagents used in this study were obtained from Sigma, USA, unless otherwise specified.

### Compounds and ionizing radiation

Antitumor, cytotoxic compounds for DNA damage induction - doxorubicin, bleomycin, phorbol 12-myristate 13-acetate (PMA) - were used at concentrations corresponding to the IC50 for specific lines. For irradiation of tumor cells with gamma photons, a radiotherapy unit RUM-17 was used, provided for work by the Military Medical Academy named after S.M. Kirov. A pre-selected therapeutic dose of 10 Gray (Gy) was used in the experiments. The irradiation parameters included a voltage across the tube of 180 kV, a current of 10 mA, a focal length of 50 cm, a 1-mm Al filter, a 0.5-mm Cu filter, and a dose rate of 0.32 Gy/min.

### Cell viability analysis

To study the effect of the bleomycin and 10Gy ionizing radiation on the cellular metabolic activity in the condition of TP53, ATOX1 and both genes inactivation, the MTT assay was used [57]. The number of surviving cells was determined by the optical density of a solution of reduced MTT (3-(4,5-dimethyl-2-thiazolyl)-2,5-diphenyl-2H-tetrazolium bromide) dye with NADP-H-dependent oxidoreductases at a wavelength of 570 nm.

### Cell cycle assay

The distribution of cell cycle (according to DNA ploidy) was analyzed on a CytoFlex B2-R2-V0 flow cytometer (USA) in PE or Rhodamine channels. A 2D PE-W versus PE-A plot was used to exclude cell conglomerates. Accumulated 20,000 events for each sample. The indicators were analyzed in the areas subG_1_, G1 and G_2_/M.

### Reverse transcription

Isolation of total RNA from cells was performed using the Total RNA isolation protocol with ExtractRNA buffer (Evrogen, Russia) according to the manufacturer’s protocol. cDNA was generated from total RNA (2 μg) by using MMLV reverse transcriptase (Evrogen, Russia). Reverse transcription PCR reaction conditions were as follows: 25°C-10 min, 42°C-50 min, 70°C-10 min, 10°C-10 sec.

### Real-time PCR analysis

For real-time PCR, a mixture for PCR was prepared, which included: a mixture of 5x qPCR SYBR Green I (Evrogen); forward and reverse primers, 10 μM each; nucleases free H2O. A negative control was prepared: a sample without the addition of the corresponding cDNA. Amplification conditions:

- Stage 1 (1 cycle): 94°C-3 min, 60°C-40 sec, 72°C-40 sec
- Stage 2 (28-30 cycles): 94°C-10s, 60°C-10s, 72°C-20s
- Stage 3 (1 cycle): 72°C-3 min
- Stage 4 (storage): 4°C

After the completion of the reactions, the expression of the products was determined by the ΔCt method, where Ct (threshold cycle) is the cycle at which the fluorescence level reaches a certain value (preselected threshold), and Δ is the change in the expression of the gene of interest relative to the reference gene, which is selected as normalization. In the experiment, transcripts of the GAPDH and HPRT genes were used for normalization. In all groups, differences from control were significant at p ≤ 0.05 (one-way ANOVA test).

The primers used are listed in Table 1.

**Table 1.**
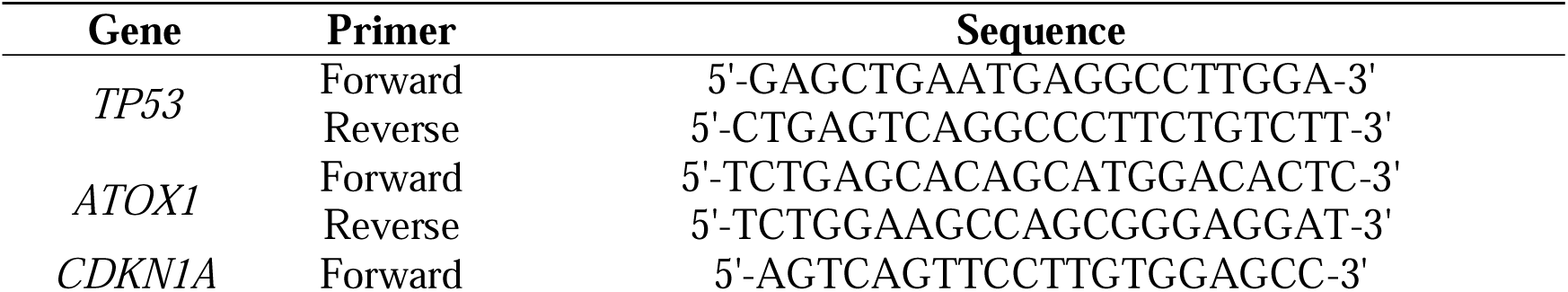
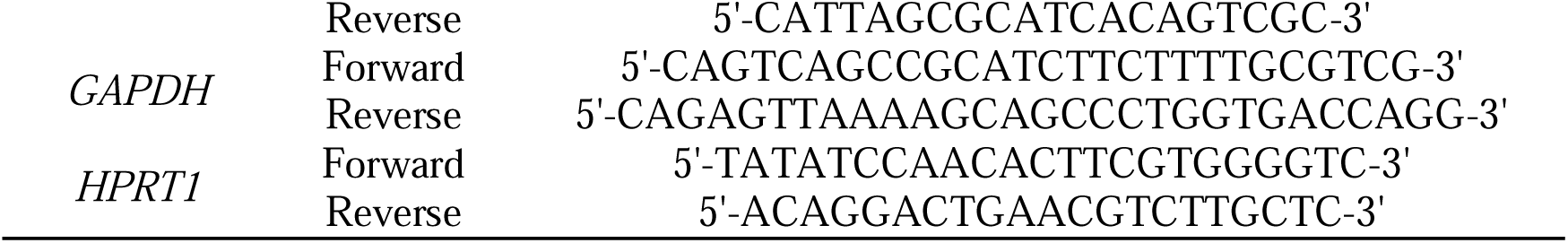
List of PCR primers.

### siRNA transfection

Lipofectamine 2000 (Invitrogen) was used to transfect siRNAs according to the manufacturer’s instructions in OptiMEM media. Transfection of siRNA was carried out 24 hours before treatment with DNA damage drugs or ionizing radiation using 250 pmol of siRNA. GFP sequences were used as scrambled RNA.

The siRNAs used are listed in Table 2.

**Table 2.**
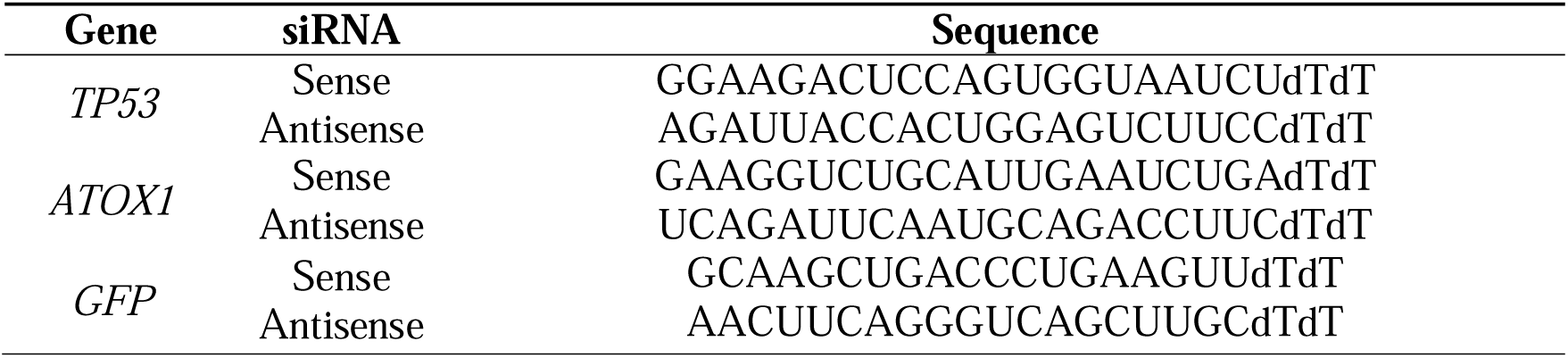
List of siRNAs.

### Western blotting

Protein electrophoresis was conducted using a polyacrylamide (PAGE) gel containing 10% SDS. A total of 35 µg of total protein was added to the gel lanes. Following electrophoresis, the proteins were transferred to a nitrocellulose membrane (Amersham, USA) using Tris-Glycine buffer. The membranes were then incubated overnight at 4°C with primary antibodies targeting p53, p21, and Atox1 proteins (Abcam, diluted 1:500-1:2000 in TBST). Anti-β-actin antibodies, diluted 1:1000, were used as an internal control. Protein visualization was achieved through chemiluminescence using secondary antibodies specific to mouse or rabbit IgG (Amersham, USA) conjugated with horseradish peroxidase. Secondary antibody dilutions ranged from 1:2000 to 1:5000. Detection was performed utilizing the ChemiDoc Touch gel-documentation system (BioRad). Densitometry analysis to evaluate the relative protein content was conducted using the Grey Mean Value Calculation tool in the ImageJ program.

### Immunofluorescence staining

Immunofluorescence staining was performed by fixing cells with 4% paraformaldehyde (PFA), permeabilizing them with 0.2% Triton X, and blocking with 1% bovine serum albumin. Cells were then incubated overnight at 4°C with primary antibodies targeting Atox1 (Abcam, diluted 1:300). Subsequently, cells were incubated with Alexa Fluor-conjugated secondary antibodies (Thermo Fisher Scientific) for 1 hour at room temperature. Nuclear staining was achieved using DAPI. Images were captured using the fluorescence microscope Leica DMi8.

### Statistical methods

Prism 8 (GraphPad) was used for statistical analysis. For the results of cell culture and immunostaining experiments, Student’s t-test was used to calculate P values. Mean ± standard deviation (SD) is shown in the figures. Mean ± standard error of the means (SEMs) is shown in the figures. Differences were considered to be significant if *p < 0*.*05*.

## Abbreviations

ROS: Reactive oxygen species
CDKN1A: cyclin-dependent kinase 1A inhibitor (Tp21)
CCND1: Cyclin D1 gene
WT: wild type cells
KO: cells without TP53 gene knockout
PMA: phorbol-12-myristate-13-acetate

## Supplementary Information

## Acknowledgments

The authors express their deep gratitude to the scientific consultant, MD Shtil Alexander for mentoring and advice on setting up experiments and interpreting the results.

## Authors’ Contributions

Conceptualization, OK and ST; investigation, AR, ST, OK; validation, AR, ST; experiments with irradiation, - VZ; original draft preparation, OK, ST, VZ; review and editing, ST, AR, OK; visualization, AR, OK; supervision and funding acquisition, OK. All authors have read and agreed to the published version of the manuscript.

## Availability of data and materials

All data generated and analyzed for this study are available from the corresponding author upon reasonable request.

## Declarations

### Ethics approval and consent to participate

Not applicable.

### Consent for publication

Not applicable.

