## Supplementary figures for "The p53 Protein is a Suppressor of Atox1 Copper Chaperon in Tumor Cells Under Genotoxic Effects"

**Supplementary materials 1.**

The online version contains supplementary material available at https://doi.org/….

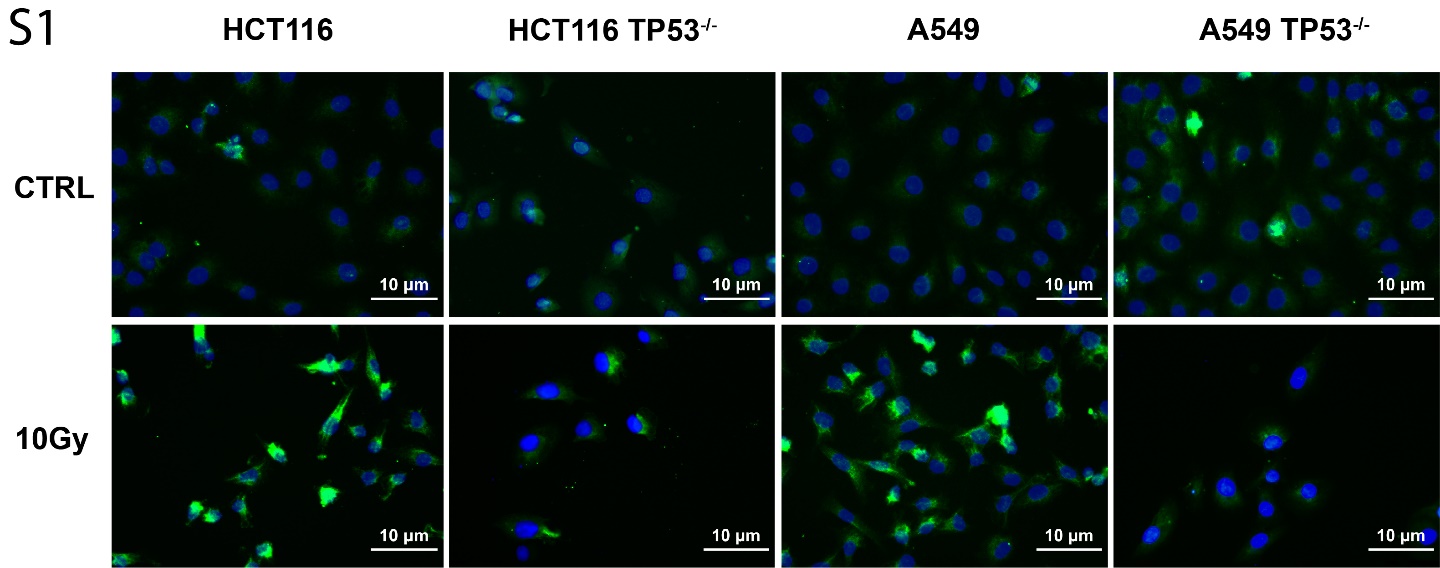

**Fig. S1.** Influence of ionizing radiation on the activity of Atox1 at different status (WT and KO) of the TP53 gene in A549 and HCT116 cell lines, 24h after ionizing irradiation (10Gy) exposure. Immunofluorescence staining with primary antibodies to Atox1, secondary antibodies with AlexaFluor488. DAPI was used for nuclei staining. TP53^-/-^ – cells without TP53. For all experiments: n = 3, mean +/− SEM, two-way ANOVA, p < 0,05.

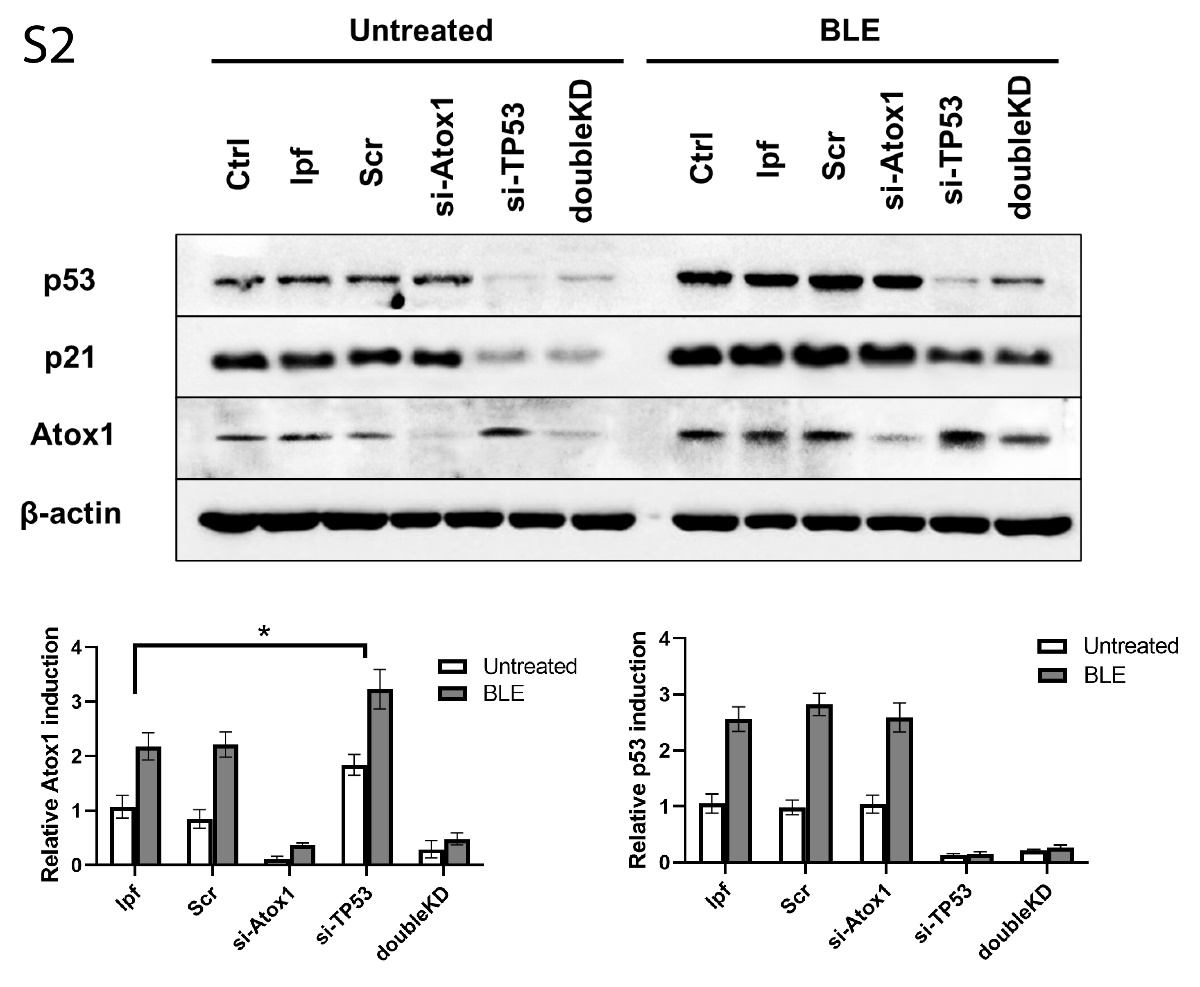

**Fig. S2.** Analyzing the effect of ATOX1 and TP53 gene siRNA-mediated knockdown on mutual expression, HCT116 cell line. Immunoblotting with antibodies to p53, p21, and Atox1; beta-actin was used as a normalization. ATOX1 (si-ATOX1), TP53 (si-TP53) or double ATOX1/ TP53 (doubleKD) knockdowns were used in the absence (Untreated) and presence (10μM BLE) of bleomycin, 24h after exposure.
